# Interference underlies attenuation upon relearning in sensorimotor adaptation

**DOI:** 10.1101/2024.05.27.596118

**Authors:** Guy Avraham, Richard B Ivry

## Abstract

Savings refers to the gain in performance upon relearning a task. In sensorimotor adaptation, savings is tested by having participants adapt to perturbed feedback and, following a washout block during which the system resets to baseline, presenting the same perturbation again. While savings has been observed with these tasks, we have shown that the contribution from implicit sensorimotor adaptation, a process that uses sensory prediction errors to recalibrate the sensorimotor map, is actually attenuated upon relearning (Avraham et al., 2021). In the present study, we test the hypothesis that this attenuation is due to interference arising from the washout block, and more generally, from experience with a different relationship between the movement and the feedback. In standard adaptation studies, removing the perturbation at the start of the washout block results in a salient error signal in the opposite direction to that observed during learning. As a starting point, we replicated the finding that implicit adaptation is attenuated following a washout period in which the feedback now signals a salient opposite error. When we eliminated visual feedback during washout, implicit adaptation was no longer attenuated upon relearning, consistent with the interference hypothesis. Next, we eliminated the salient error during washout by gradually decreasing the perturbation, creating a scenario in which the perceived errors fell within the range associated with motor noise. Nonetheless, attenuation was still prominent. Inspired by this observation, we tested participants with an extended experience with veridical feedback during an initial baseline phase and found that this was sufficient to cause robust attenuation of implicit adaptation during the first exposure to the perturbation. This effect was context-specific: It did not generalize to movements that were not associated with the interfering feedback. Taken together, these results show that the implicit sensorimotor adaptation system is highly sensitive to memory interference from a recent experience with a discrepant action-outcome contingency.

## Introduction

People exhibit a remarkable ability to retain motor skills. The skier finds herself performing close to her previous level of competence after an 8-month layoff and, while the mind might be wary, our bodies are able to readily recall how to ride a bike even if they have not done so for many years. Indeed, laboratory studies of retention have highlighted the phenomenon of savings, the benefit in performance observed when relearning a previously acquired skill. Originally described in the verbal learning domain (Ebbinghaus, 1913), savings has since come to be recognized as a rather ubiquitous phenomenon, including many examples in the motor learning literature (Coltman et al., 2019; Doyon et al., 2009; Kojima et al., 2004; Mawase et al., 2014; Medina et al., 2001; Milner, 1962; Walker et al., 2002).

Sensorimotor adaptation tasks have provided an interesting testbed for exploring savings in motor learning. In visuomotor adaptation, feedback indicating the participant’s hand position is perturbed, usually by imposing an angular rotation. Participants learn to compensate, reaching in the opposite direction of the perturbation. Savings is tested by presenting the same perturbation over two learning blocks, separated by a washout phase with non-perturbed feedback (Albert et al., 2022; Herzfeld et al., 2014; Huang et al., 2011; Krakauer, 2009; Krakauer et al., 2005; Zarahn et al., 2008). With this design, participants often exhibit savings, showing a higher rate of adaptation during the second learning block.

In line with the common conception about the procedural nature of motor skills, this form of savings has been hypothesized to reflect changes in an implicit system that serves to keep the sensorimotor system well-calibrated (Albert et al., 2022, 2021; Coltman et al., 2019; Hadjiosif et al., 2023; Yin and Wei, 2020). For example, it has been proposed that the sensitivity of this system is enhanced when participants are exposed to errors during initial learning, resulting in faster learning upon re-exposure to similar errors (Albert et al., 2021; Herzfeld et al., 2014).

However, by using methods that isolate the contribution of this recalibration process, we have found that implicit sensorimotor adaptation is actually attenuated during relearning (Avraham et al., 2021; see also Hamel et al., 2022, 2021; Leow et al., 2020; Stark-Inbar et al., 2016; Tsay et al., 2021b; Wang and Ivry, 2023; Yin and Wei, 2020). Savings, when observed in adaptation tasks, appears to largely reflect the operation of other learning processes such as the recall of a successful explicit strategy (Avraham et al., 2021; Haith et al., 2015; Huberdeau et al., 2019; Morehead et al., 2015).

Why does implicit sensorimotor adaptation show attenuated relearning rather than savings? We can consider two hypotheses. The first hypothesis is also based on the idea that exposure to a given error changes the sensitivity of the system in response to that error when re-encountered. However, rather than increasing with familiarity, sensitivity may decrease with familiarity. That is, the adaptation system becomes less responsive after prolonged exposure to an error. Mechanistically, desensitization could arise from a saturation of recently activated synapses (Hamel et al., 2022, 2021; Nguyen-Vu et al., 2017).

An alternative hypothesis is based on the idea of interference, a classical concept in the psychological memory literature (Underwood, 1957). In the typical design of a savings study, the first block with perturbed feedback is followed by a washout block with veridical feedback to reset the behavior back to baseline. When exposed to veridical feedback in the adapted state, the participant experiences an error in the opposite direction of that experienced during the initial learning block. Thus, at the end of the washout block, the participant has been exposed to two error signals of opposite signs, one associated with the initial perturbation and a second associated with the washout. When the perturbation is re-introduced, memories associated with these two error signals may compete with one another (Bouton, 1986), especially given that the context is similar (reaching to the same target). As such, the response to the re-introduced perturbation may be attenuated due to interference from the association established during the washout block.

We evaluate these two hypotheses in a series of behavioral experiments.

## Results

We employed a visuomotor task in which task-irrelevant clamped feedback was used to isolate implicit adaptation (Morehead et al., 2017) (Fig. 1A). In this task, participants reach to a visual target while the cursor follows a path with a fixed angular offset from the target (e.g., by 15°). Despite instructions to ignore the cursor, participants show robust implicit adaptation to this type of perturbation, with the direction of the hand movement (reach angle) gradually shifting away from the target (and cursor). In our previous study, we used this manipulation over two learning blocks that were separated by a washout block and found that the implicit change in reach angle during the second learning block was attenuated with respect to the first learning block (Avraham et al., 2021).

**Fig 1.**
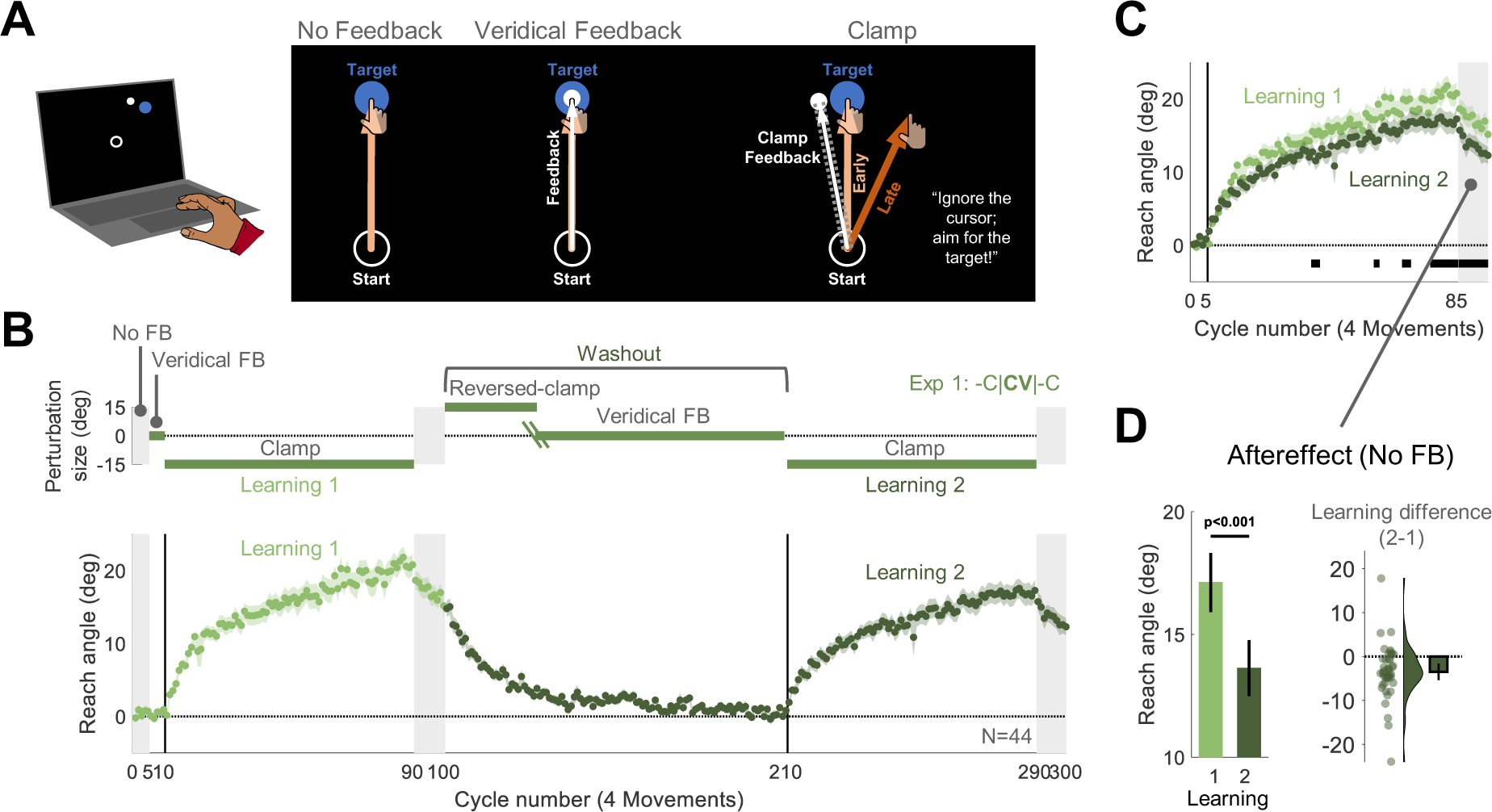
Experiment 1: Upon relearning a visuomotor rotation, implicit adaptation is attenuated. (**A**) Task design and schematics of all trial types in Experiment 1. Using a trackpad or mouse, participants (N=44) moved a cursor from the start location (white circle) to a target (blue disk), with the target appearing at one of four locations (one representative location is depicted). There were 3 types of trials: 1) No feedback, with the cursor disappearing at movement onset; 2) Veridical feedback, in which the direction of the cursor (small white disk) was veridical with respect to the direction of the movement; 3) Clamped feedback, in which the cursor followed an invariant path with respect to the target. (**B**) Top: Experimental protocol. The -C|**CV**|-C abbreviation indicates the main block-level structure of the experiment. There were two learning blocks with clamped feedback (-C), each followed by an aftereffect block with no feedback. To reset the sensorimotor map following the first learning block, we used a washout block composed of a reversed-clamp feedback phase (C) and a veridical feedback phase (V). The green oblique lines in the washout block mark the transition between the two phases, with the length of the reversed clamp phase determined on an individual basis (see Methods). Bottom: Time course of mean reach angle, with the data averaged within each cycle of four movements. Light and dark shading signify learning blocks 1 and 2, respectively, with the onset of the clamped feedback marked by the vertical solid lines. (**C**) Overlaid reach angle functions for the two learning blocks and two aftereffect blocks. Horizontal thick black lines denote clusters that show a significant difference between blocks 1 and 2 (*p* < 0.05). (**D**) The left panel (pair of bars) presents the aftereffect data (mean ± SEM) for each learning block, measured as the averaged reach angle across all cycles in each aftereffect block. The right panel shows the within-participant difference (Aftereffect 2 – Aftereffect 1; dots and violin plot-distribution of individual difference scores, bar-mean difference and 95% CI). SEM, standard error of the mean. CI, Confidence Interval.

In Experiment 1, we aimed to replicate this attenuation effect using a similar relearning protocol (abbreviated as -C|**CV**|-C, Fig. 1B). There were two learning blocks with -15° clamped feedback (-C), separated by a washout block. The washout block started with reversed clamped feedback, with the cursor now shifted in the opposite direction (15°, **C**). This was expected to implicitly drive the adapted hand movements back towards the target. Once the participant’s reaches were near the target, we transitioned to veridical feedback (V) for the remainder of the washout block. In this manner, the average reach angle at the start of the two clamp blocks should be similar. We also included a short no-feedback block after each learning block to assess the aftereffect. During all phases of the experiment, the instructions remained the same: Reach directly to the target.

Participants exhibited robust adaptation during the first learning block, and their behavior reverted back to baseline during the washout block. Consistent with our previous findings, the overall implicit change in reach angle during the second learning block was smaller than during the first learning block (Figs. 1B, 1C). To evaluate this effect statistically, we compared the reach angle between the first and second learning blocks, averaging the data over cycles of four reaches (one reach per target). We used a non-parametric cluster-based permutation test to identify clusters of cycles that differed between the two learning blocks (Avraham et al., 2021; Labruna et al., 2019; Maris and Oostenveld, 2007). This analysis revealed multiple clusters in which there was a decrease in reach angle during the second learning block. In line with the cluster-based results, attenuation was also significant when the analysis was restricted to the aftereffect measure of adaptation, namely the mean reach angle during the aftereffect blocks ([mean difference, 95% CI], −3.48°, [-5.36° -1.61°], t(43) = −3.75, *p* = 5.20×10^-4^, BF10 = 53.2, d = −0.57, Fig. 1E).

In Experiment 1, relearning was tested after a washout block in which participants had performed 110 reaches to each target in the presence of feedback – either reversed-clamp or veridical – that was different than the error signal that drove initial learning. It is unclear if the attenuation is due to the exposure to these alternative and potentially interfering forms of feedback or reflects desensitization to the original error. Experiment 2 was designed to address this question. As in Experiment 1, participants experienced two learning blocks that were separated by a washout block (Fig. 2A). However, no feedback was presented during the entire 110-cycle washout block (-C|**N**|-C design). If attenuation upon relearning is due to interference from the feedback experienced during washout, removing this feedback should rescue attenuated learning. In contrast, if attenuation is driven by desensitization to a familiar error, we should again observe attenuation in the second learning block following the no-feedback washout block.

**Fig 2.**
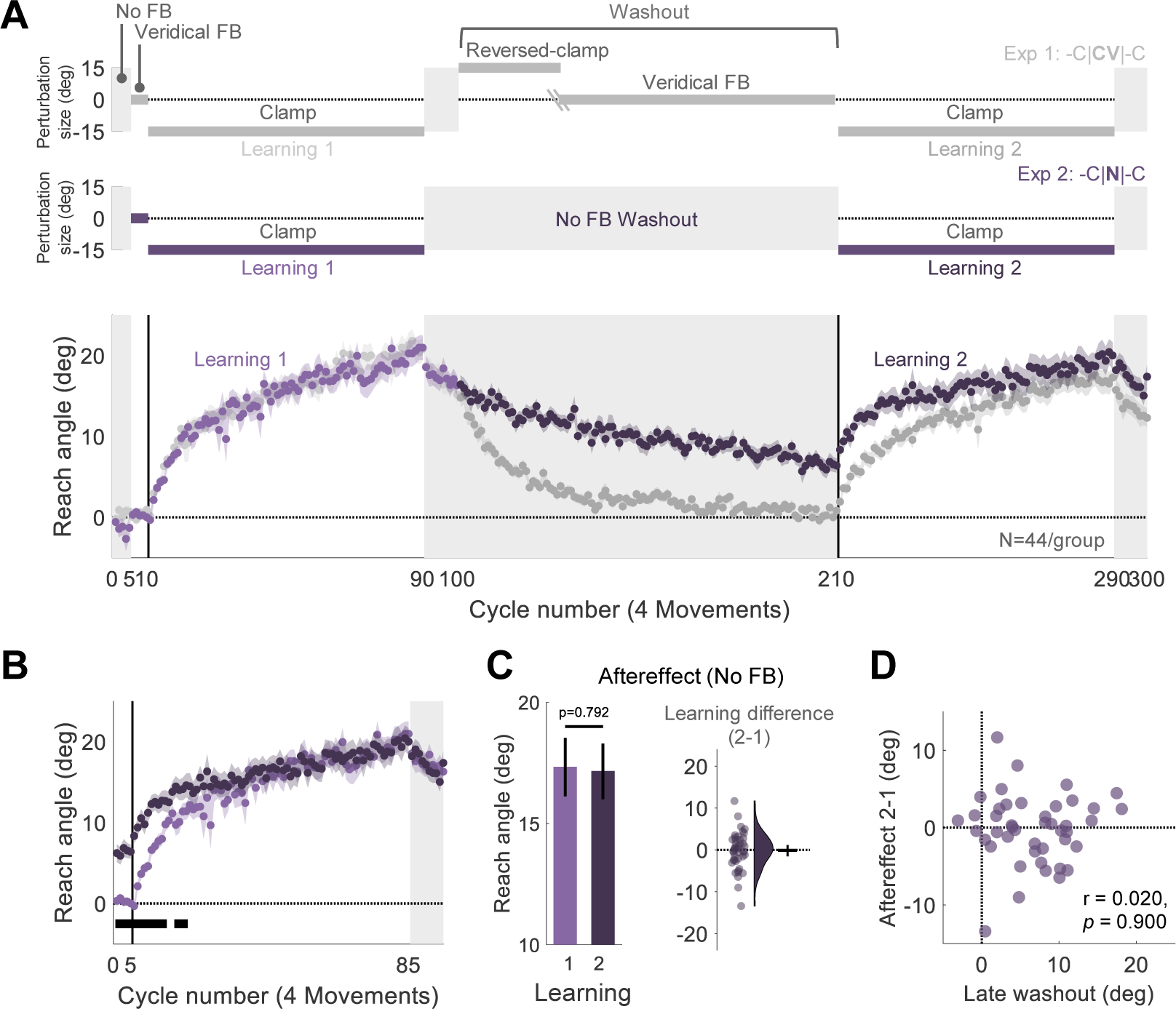
Experiment 2: Implicit adaptation is not attenuated upon relearning when feedback is eliminated during the washout block. (**A**) Experimental protocol and learning functions. Top: Participants (N=44) experienced 110 cycles of trials without feedback (N) during a washout block that separated the two learning blocks (-C|**N**|-C design, purple). Bottom: Time course of mean reach angle averaged over cycles. Light and dark shading signify learning blocks 1 and 2, respectively, with the onset of the clamped feedback marked by the vertical solid lines. The design and learning functions from Experiment 1 are reproduced here to provide a visual point of comparison (gray). (**B**) Overlaid reach angle functions for the two learning blocks and two aftereffect blocks in Experiment 2 (significant clusters based on p<0.05) (**C**) The left panel presents the aftereffect (mean ± SEM) for each learning block and the right panel the within-participant difference (Aftereffect 2 – Aftereffect 1; dots and violin plot-distribution of individual difference scores, bar-mean and 95% CI). (**D**) Scatter plot showing no relationship between the reach angle at late washout and change in relearning (Aftereffect 2 – Aftereffect 1). SEM, standard error of the mean. CI, Confidence Interval.

Despite the absence of feedback, adaptation slowly decayed throughout the washout block, presumably due to some form of forgetting or attraction to the original state of the sensorimotor system (Fig. 2A). The decay was, on average, incomplete by the end of the washout block. Thus, the mean reach angle at the beginning of the second learning block was higher than at the start of the first Learning block (Fig 2B). However, after ∼20 cycles, the two functions converged and reached a similar state of adaptation by the end of the clamp blocks. Thus, we failed to observe attenuation. This null result was confirmed in the analysis of the aftereffect data (−0.18°, [-1.52° 1.16°], t(43) = −0.266, *p* = 0.792, BF10 = 0.169, d = −0.04, Fig. 2C). Interestingly, the difference between reach angle in the first and second aftereffect blocks was unrelated to the extent of decay observed during the washout block (r = 0.020, *p* = 0.900, BF10 = 0.12). As can be seen in Figure 2D, even those participants who showed complete washout in the absence of feedback did not exhibit attenuated relearning.

In summary, the results of Experiment 2 indicate that attenuation upon relearning is not due to desensitization to the error that participants experienced during initial learning. Rather, the results are consistent with the hypothesis that attenuation is due to interference that arises from the feedback experienced during the washout block.

What conditions might produce interference during relearning? With the introduction of the reversed clamp at the beginning of the washout block in Experiment 1, participants are exposed to a large error signal that is in the opposite direction to that experienced during the initial learning block. Interference may arise from a competition between the memory traces associated with the two opposing error signals (Heald et al., 2021; Sing and Smith, 2010). This hypothesis can account for the results of Experiment 2 since there was no error presented during the washout block. An even stronger test would be to use feedback during the washout block that did not provide salient error information that was opposite to that observed during the initial learning block.

We implemented this test in Experiment 3. Following implicit adaptation with the rotated clamped feedback, participants were told at the start of the washout block that the cursor would now be aligned with their hand. However, we actually applied a visuomotor rotation that was contingent on the movement direction of the hand, setting the size of the rotation so that the cursor would move close to the target (see Methods). The size was determined on an individual basis: For example, if the participant had adapted 19° in the clockwise direction at the end of the rotation block, the rotation was initially set to 19° in the counterclockwise direction. This perturbation was gradually reduced over the washout block (Fig. 3A). Gradual changes in the magnitude of a visuomotor rotation have been shown to result in pronounced implicit adaptation (Christou et al., 2016; Modchalingam et al., 2023). Here, we used this method to nullify an adapted state. When the size of the rotation had decreased to 0°, the feedback was veridical and remained so for the rest of the washout block. We assumed that this method would minimize the participant’s experience with salient opposite-signed errors during the early (and all) stages of the washout block despite having adapted during the initial learning block.

**Fig 3.**
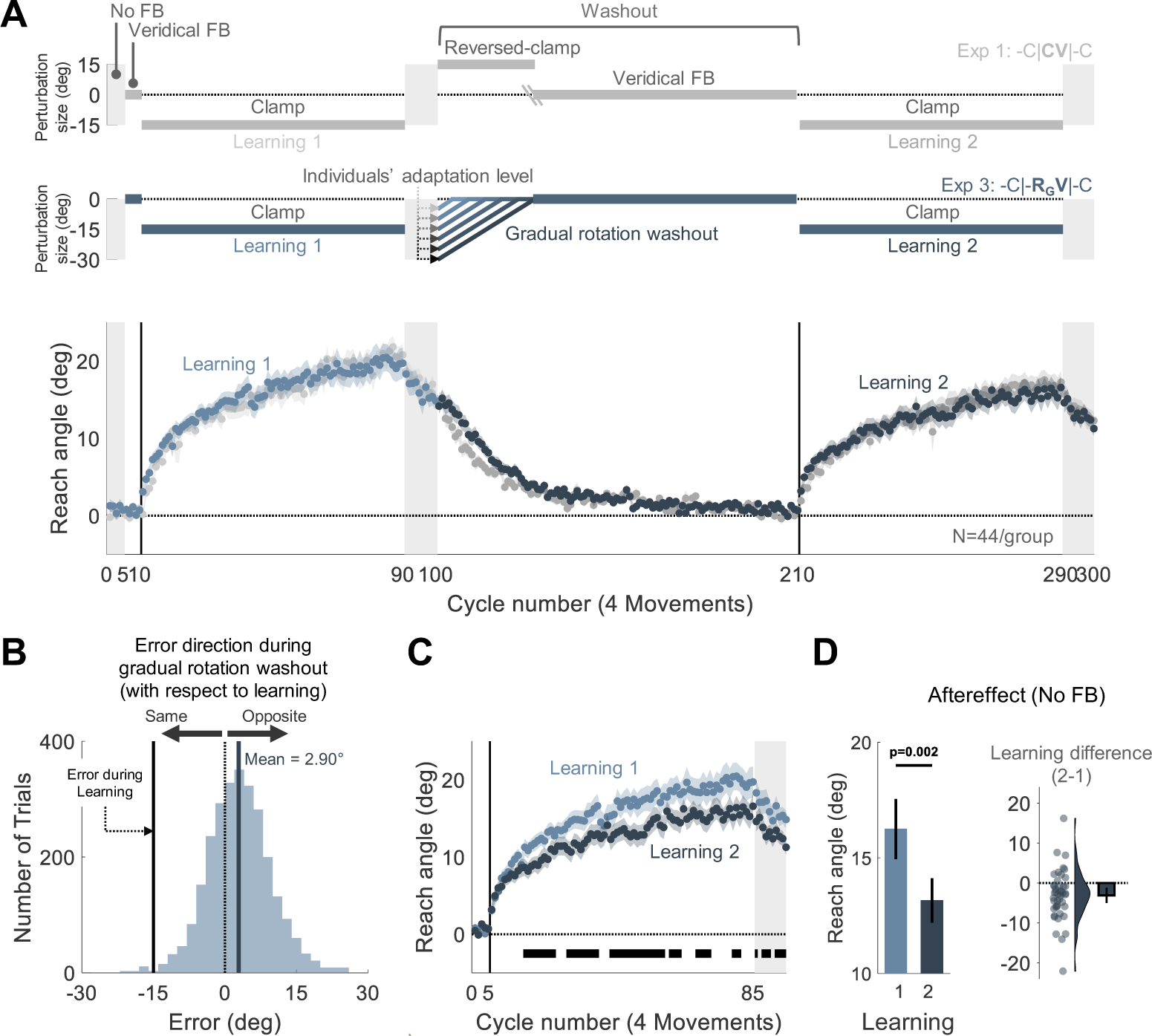
Experiment 3: Attenuated adaptation does not require experience with salient, opposite signed error at the beginning of washout. (**A**) Experimental protocol and learning functions. Top: At the beginning of the washout block in Experiment 3, participants (N=44) experienced a rotated cursor that was contingent on the direction of their hand movement, with the magnitude of the perturbation set to their final adaptation level at the end of the first learning block; in this way, the cursor position would be near the target. The size of the rotation was gradually decreased until reaching 0°, at which point it was veridical and remained so for the rest of the washout block (-C|-**RGV**|-C design, blue). Bottom: Time course of mean reach angle averaged over cycles (4 movements). Light and dark shading signify learning blocks 1 and 2, respectively, with the onset of the clamped feedback marked by the solid vertical lines. The design and learning functions from Experiment 1 are reproduced here to provide a visual point of comparison (gray). (**B**) Distribution of errors experienced during the non-zero rotation phase of the washout block. These errors were small in magnitude (mean=2.9°, dark blue solid line) and in the opposite direction of the error experienced during the initial learning block (black solid line). Presumably, these opposite errors are the signals that drive the washout of the initial adaptation. The dotted line represents zero error. (**C**) Overlaid reach angle functions for the two learning blocks. Horizontal thick black lines denote clusters that show a significant difference between blocks 1 and 2 (*p* < 0.05). (**D**) The left panel presents the aftereffect (mean ± SEM) for each learning block and the right panel the within-participant difference (Aftereffect 2 – Aftereffect 1; dots and violin plot-distribution of individual difference scores, bar-mean and 95% CI). SEM, standard error of the mean. CI, Confidence Interval.

After adaptation, participants exhibited a gradual reversion of reach angle back to baseline (Fig. 3A). This washout function was somewhat slower than that observed in Experiment 1 where we had used a salient, reversed clamp to achieve washout (Fig. S1), presumably because the experienced errors during the initial washout trials were much smaller and of variable sign in Experiment 3 ([mean ± standard deviation], 2.90° ± 10.1°, Fig. 3B) (Avraham et al., 2020; Kim et al., 2018; Tsay et al., 2021a; Wei and Körding, 2009).

Contrary to our prediction, the elimination of salient, opposite-signed errors did not abolish the attenuation effect. Robust attenuation was observed across the second learning block and during the aftereffect block ([mean aftereffect difference, 95% CI], −3.09°, [-4.99° - 1.18°], t(43) = −3.27, *p* = 0.002, BF10 = 15.11, d = −0.49, Figs. 3C, 3D).

While the results indicate that attenuation does not require a salient change in the error, an analysis of the feedback provided at the beginning of the washout block shows that participants experienced opposite-signed errors more often than same-signed errors (Fig 3B. Fig S2 provides a comparison with the error distribution experienced with veridical feedback). The small skew in the distribution might be sufficient to produce interference.

Alternatively, interference may arise when multiple associations are linked to the same action. Up to this point, we had assumed that interference would occur when a given action (e.g., moving to a given target location) was associated with two types of perturbations, specifically, opposing visuomotor errors. However, simply experiencing a change in the feedback, even from one that is not perturbed, may be sufficient to elicit interference.

We tested this hypothesis in Experiment 4 using a protracted block with veridical feedback, asking if this would attenuate subsequent adaptation to a perturbation. On each trial, the target appeared within one of two 30°-wide wedges located at opposite poles of the workspace (∼180° separation, Fig. 4A). When the target appeared in one of the wedges, the perturbation followed a similar schedule to Experiment 1: Two learning blocks with clamped feedback were separated by a washout block -C|**CV**|-C). When the target appeared in the other wedge, the feedback was veridical for the first 85 cycles, followed by a single learning block with clamped feedback (**V85**|-C). Previous work has shown negligible generalization of adaptation for locations separated by more than 45° (Krakauer et al., 2000; McDougle et al., 2017; Morehead et al., 2017). As such, this within-participant design allowed us to ask if extended exposure to veridical feedback was sufficient to attenuate adaptation. Specifically, we asked if adaptation to targets in the **V85**|-C location would be attenuated with respect to the first learning block in the - C|**CV**|-C location.

**Fig 4.**
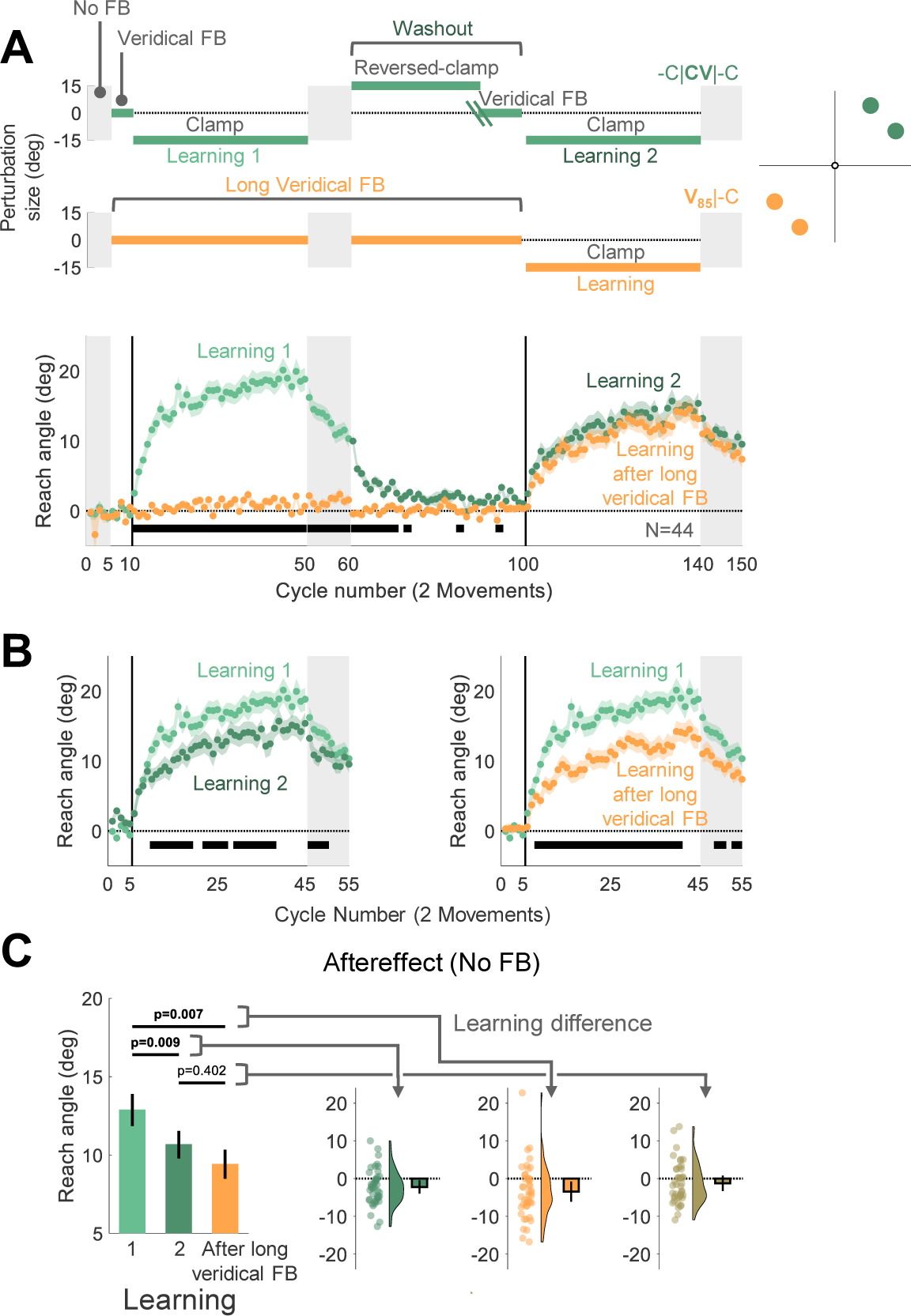
Experiment 4: Adaptation is attenuated for movements that were previously associated with either washout after learning or an extended baseline experience with veridical feedback. (**A**) Experimental protocol and learning functions. Top: The target appeared at one of four locations, with two locations falling within one 30°-wedge and the other two falling within a same-size wedge on the opposite side of the workspace. Participants (N=44) experienced a - C|**CV**|-C schedule (cyan) in one wedge and a **V85**|-C design schedule in the other wedge (orange). For the latter, the number of veridical feedback cycles (85) matched the total number of cycles before relearning at the other wedge (excluding the no-feedback trials). Bottom: Time course of mean reach angle averaged over cycles, with each cycle consisting of 2 movements for each wedge. Light and dark cyan signify learning blocks 1 and 2 in the -C|**CV**|-C condition. (B) Overlaid reach angle functions comparing the first learning block in -C|**CV**|-C to that of the second learning block in the same wedge (left panel) and to the post long-baseline learning block at the other wedge (right panel). Horizontal thick black lines (B) and (C) denote clusters that show a significant difference between the functions (with *p* < 0.017 as a significance criterion, see Methods). (**C**) Left panel (bars) presents the aftereffect data (mean ± SEM) for each learning block and the right panel shows the within-participant differences for three contrasts: 1) Aftereffect 2 – Aftereffect 1; 2) Aftereffect after long baseline – Aftereffect 1; 3) Aftereffect after long baseline – Aftereffect 2. Dots and violin plots show the distribution of individual difference scores; bar-mean and 95% CI. SEM, standard error of the mean. CI, Confidence Interval.

For the -C|**CV**|-C location, adaptation in the second learning block was attenuated with respect to the first learning block, consistent with the results of Experiment 1 (Fig. 4A, 4B). Critically, adaptation in the **V85**|-C location was also attenuated relative to the first learning block in the -C|**CV**|-C location, and it was comparable to the second learning block in the -C|**CV**|-C location. This pattern was also evident in the aftereffect data (F[1.59, 68.5] = 7.96, *p* = 0.002, BF10=40.95, and η ^2^=0.16, mean difference Aftereffect 2 – Aftereffect 1: -2.21°, 95% CI: [-3.97° - 0.45°], *pB* {Bonferroni corrected *p* value} = 0.009, mean difference Aftereffect after long baseline – Aftereffect 1: -3.46°, 95% CI: [-6.14° -0.79°], *pB* = 0.007, mean difference Aftereffect after long baseline – Aftereffect 2: -1.25°, 95% CI: [-3.29° 0.79°], *pB* = 0.402, Fig. 4C). Thus, attenuation of implicit adaptation does not require experience with an opposing perturbation; rather, it can also arise following extended experience with veridical feedback.

Experiment 4 was designed based on the assumption that the mechanisms underlying attenuation do not generalize across the workplace. Although it is well-established that implicit adaptation shows minimal generalization between regions separated by more than 45° (Krakauer et al., 2000; McDougle et al., 2017; Morehead et al., 2017), generalization rules may be different for attenuation. For example, if attenuation generalizes broadly, the reduced rate of learning following extended exposure to veridical feedback at one wedge could be due to interference arising from exposure to the reversed clamp at the other wedge.

Given this concern, we used an alternative design in Experiment 5 to assess the effect of extended experience with veridical feedback on attenuation. The target appeared in one of three wedges, with each wedge separated by at least 90° from the other two (Fig. 5A). In all three wedges, there was a single learning block with clamped feedback, with the direction of the perturbation the same across wedges. We varied the number of veridical feedback cycles that preceded the clamped feedback block, using 5 cycles for one wedge (**V5**|-C), 45 cycles for a second wedge (**V45**|-C), and 85 cycles for the third wedge (**V85**|-C). If experience with veridical feedback is sufficient to cause interference in a context-specific manner, we expect that the level of adaptation will be inversely related to the number of veridical cycles.

**Fig 5.**
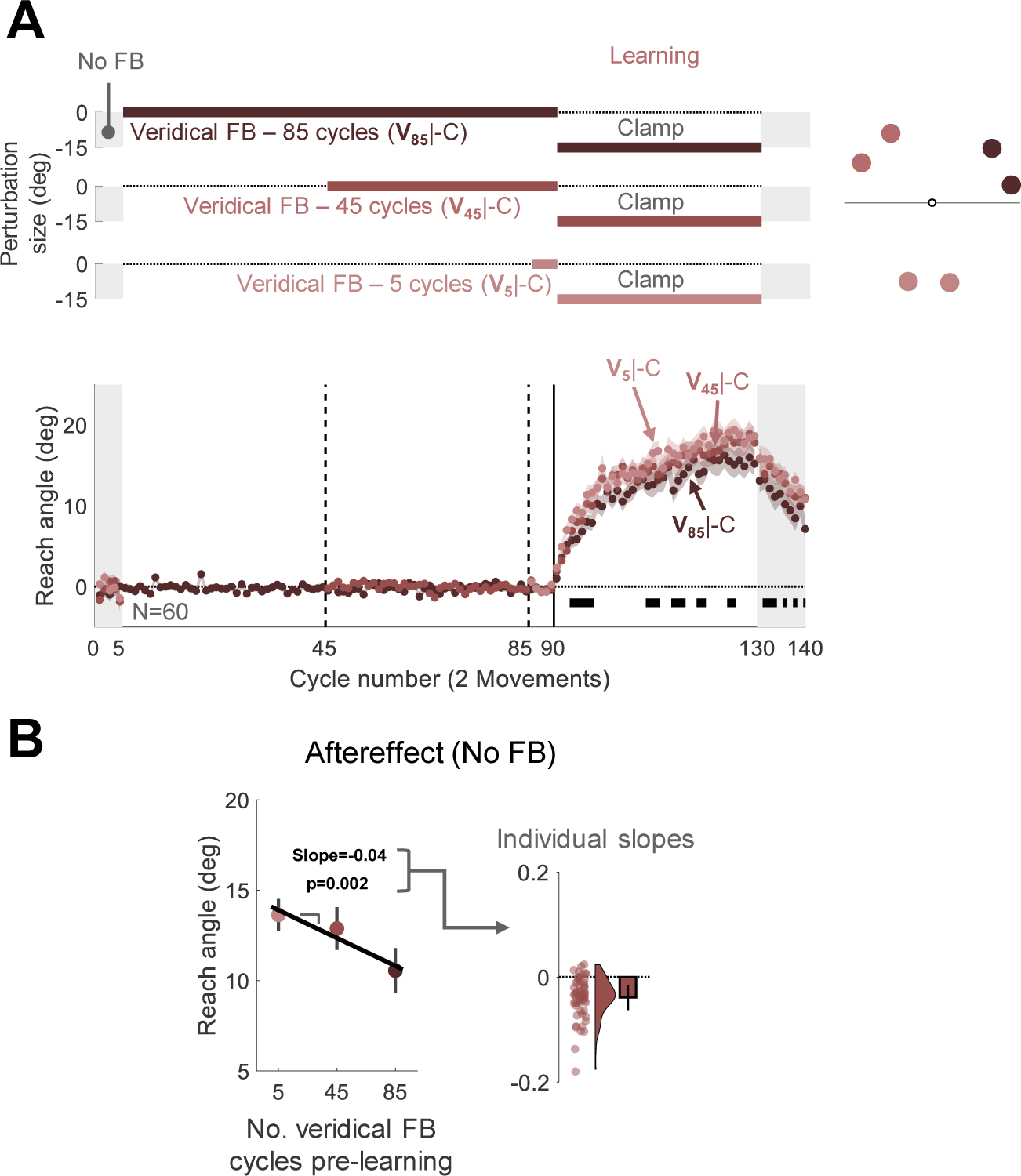
Experiment 5: Interference from veridical feedback is context-specific. (**A**) Experimental protocol and learning functions. Top: During the learning block, participants (N=60) experienced rotated clamped feedback while reaching to six targets, with two targets falling within each of three wedges distributed across the workspace. For each wedge, participants experienced a different number of cycles with veridical feedback prior to the learning block: 5 (**V5**|-C, light red); 45 (**V45**|-C, medium red); 85 (**V85**|-C, dark red). Bottom: Time course of mean reach angle averaged over cycles (2 movements) for each wedge. The vertical solid lines at cycles 45 and 85 mark the onset of movements to an additional location, and the vertical solid lines at cycles 90 mark the onset of the task-irrelevant clamped feedback. Horizontal thick black lines denote clusters of cycles that show a significant relationship between the reach angle and the number of veridical cycles (*p* < 0.05). (**B**) Left panel presents the aftereffect results (mean ± SEM) for each learning condition with the fixed effect regression line obtained using a linear mixed model. Right panel shows the distribution (dots and violin plots) of individuals’ slopes (random effect); bar-mean slope and 95% CI. SEM, standard error of the mean. CI, Confidence Interval.

The results were consistent with this prediction. A nonparametric permutation test identified significant cycles during the learning and aftereffect blocks for which the reach angle was negatively associated with the number of veridical cycles (Fig. 5A). This was also evident in a linear mixed-effect analysis assessing the change in aftereffect across the three conditions (Fixed effect slope: −0.04, [-0.06 -0.01], t(178) = −3.14, *p* = 0.002, Fig. 5B). Overall, the results of Experiment 5 provide strong evidence that extended experience with veridical feedback can interfere with implicit adaptation in a context-specific manner.

## Discussion

Relearning paradigms have offered an important approach to studying learning and memory. While many studies have demonstrated savings, indicative of latent retention of information that was acquired in the past, implicit sensorimotor adaptation exhibits attenuation upon re-exposure to a previously learned perturbation (Avraham et al., 2021; Hamel et al., 2022, 2021; Leow et al., 2020; Stark-Inbar et al., 2016; Tsay et al., 2021b; Yin and Wei, 2020). The goal of this paper was to identify the mechanisms underlying this attenuation. The results indicate that attenuation is observed when an action has been associated with discrepant forms of visual feedback, creating interference between two sensorimotor memories.

### Implicit adaptation is subject to interference from discrepant feedback signals

In the adaptation literature, studies of relearning have typically used a design in which a washout, perturbation-free block is employed between two learning blocks (Herzfeld et al., 2014; Huang et al., 2011; Leow et al., 2012). The purpose of the washout block is to bring behavior back to baseline (i.e., the hand movement is directed to the visual target) before the re-introduction of the perturbation. As such, one can compare two learning functions that start, at least superficially, from a common state. When considering overall performance, savings is typically observed: Learning in the second perturbation block is faster than during the initial perturbation block.

However, the observed behavior likely reflects the operation of multiple learning processes. In particular, in addition to an implicit recalibration process, participants may employ an explicit strategy to compensate for the perturbation (Bond and Taylor, 2015; Kim et al., 2021; Taylor et al., 2014). Savings upon re-exposure to a perturbation appears to reflect the recall of a strategy that had proven to be successful in the initial perturbation block (Avraham et al., 2021; Haith et al., 2015; Morehead et al., 2015). By using non-contingent, clamped feedback, we were able to eliminate strategy use and thus, by inference, isolate learning to just that produced by implicit adaptation (Morehead et al., 2017; Tsay et al., 2020b). Under such conditions, relearning is attenuated rather than facilitated. Critically, this attenuation has also been observed in tasks that used other methods to isolate implicit sensorimotor adaptation (Avraham et al., 2021; Hamel et al., 2021; Huberdeau et al., 2019; Leow et al., 2020; Yin and Wei, 2020).

Our initial hypothesis was that this attenuation reflects interference that arises when a movement is associated with conflicting error signals. When veridical feedback is re-introduced in the washout block, a salient error is experienced, one that is in the opposite direction of that experienced during the initial perturbation block. Similarly, conflicting error signals are associated with the same movement when reversed clamped feedback is used to drive washout. Consistent with the interference hypothesis, relearning was not attenuated when feedback was omitted from the washout block. However, subsequent experiments showed that interference does not require conflicting error signals: We also observed attenuation when we used a procedure to wash out adaptation in the absence of a salient opposite error signal and, more dramatically, during an initial perturbation block that followed an extended baseline block with veridical feedback. Thus, interference appears to be between discrepant feedback signals and not competing error signals.

The attenuating effect from a prolonged baseline period with veridical feedback also helps clarify that the effect is due to anterograde interference rather than retrograde interference (see also Hinder et al., 2007; Krakauer et al., 2005; Miall et al., 2004; Villalta et al., 2015). Anterograde interference refers to the disruptive effect of one memory on the ability to either acquire or recall another memory. In contrast, retrograde interference refers to the disruptive effect of a new memory on the consolidation of a previously learned memory. The fact that attenuation is seen in the initial exposure to a perturbation is consistent with an anterograde interference account. We recognize that retrograde interference may also be operative in the experiments that included two learning blocks (Experiments 1-4). To assess this, future experiments with clamped feedback could adopt classic manipulations used to evaluate the relative contribution of anterograde and retrograde interference such as varying the delay between the initial learning block, washout block, and re-learning block. These manipulations would also be important to ask how interference effects are modulated when there is greater time for consolidation of the memory associated with the initial perturbation (Bock et al., 2001; Brashers-Krug et al., 1996; Caithness et al., 2004; Criscimagna-Hemminger and Shadmehr, 2008; Goedert and Willingham, 2002; Krakauer et al., 1999; Shadmehr and Holcomb, 1999, 1997).

### Interference or desensitization?

We have proposed that the difference observed between the two learning blocks reflects the interference of two competing memories, one associated with the feedback experienced during learning and another with the feedback experienced during washout (or from a long baseline block). By this view, the output of the system reflects the contribution from two memories, yielding a change in behavior even if the parameters of the system (e.g., learning rate) remain invariant.

An alternative perspective centers on the idea that the adaptation system retains a memory of error history and uses this information to adjust its sensitivity to the error. For example, in the Memory of Error model, the system upregulates the learning rate in response to experienced or stable errors and downregulates the learning rate in response to unfamiliar or variable errors (Albert et al., 2022, 2021; Coltman et al., 2019; Herzfeld et al., 2014; Yin and Wei, 2020). The failure to observe savings in the implicit adaptation system, together with the findings that the implicit response is the same for stable or variable errors, support the notion that an increase in learning rate (i.e., savings) is likely a property of explicit processes (Avraham et al., 2020; Hutter and Taylor, 2018; Wang et al., 2024a).

Nonetheless, attenuation upon relearning could be taken to reflect a flexible implicit system, one in which the sensitivity to error decreases over learning sessions rather than increases. For example, learning might become weaker due to the saturation or depletion of neuroplasticity mechanisms (Hamel et al., 2022, 2021). Consistent with this hypothesis, a recent study showed that attenuation was observed when there were 320 trials in the first perturbation block, but not when this number was reduced to 40 trials (De La Fontaine et al., 2023). However, to match the overall number of trials between the two conditions, the authors extended the baseline block for participants in the short learning condition. As shown in Experiments 4 and 5 above, an extended baseline block with veridical feedback attenuates learning in the first block. Indeed, in De La Fontaine’s study, the aftereffect of the first perturbation block was smaller for the long baseline/short learning group compared to the short baseline/long learning group. Thus, the absence of attenuation for the former group may be due to interference in both learning blocks.

Nonetheless, the interference and desensitization hypotheses are not mutually exclusive. Attenuation in the second learning block could be driven by a contribution from both interference and desensitization (Hamel et al., 2022, 2021). However, attenuation is neither observed when visual feedback is eliminated in the washout block (Experiment 2) nor during a prolonged perturbation block with the same error (Avraham et al., 2021; Kim et al., 2018; Morehead and Smith, 2017). Both results would appear at odds with a desensitization account, at least one in which desensitization occurs as a function of the amount of experience or time responding to a given perturbation. At present, we assume that interference is the primary mechanism underlying attenuation upon relearning, at least in visuomotor adaptation.

Hadjiosif et al. (Hadjiosif et al., 2023) have suggested that the behavior during relearning depends on the time-dependent decay characteristics of distinct memory components. One component is a temporally volatile process that can rapidly respond to a perturbation but results in a change in state that is quickly forgotten; this system is hypothesized to show heightened sensitivity to experienced errors and thus drive savings. The second component is a temporally stable process that exhibits slow learning but shows high retention over time; this system is hypothesized to exhibit attenuation upon relearning. Our results are inconsistent with the view that attenuation is restricted to a temporally stable implicit process. Given the number of target locations and short inter-trial intervals in our experiments, and based on the parameters in Hadjiosif et al.’s model, a temporally volatile process would still be making a significant contribution to behavior in some of our conditions (e.g., in Experiment 4, the median time between reaches to a given target = 1.92 s, interquartile range: [1.65 s 2.33 s], corresponding to a decay of only ∼5% (Hadjiosif et al., 2014)). Thus, we should have observed savings rather than attenuation. We hypothesize that given the methods used in their study, the behavioral changes they attribute to an implicit temporally volatile process are actually due to strategic adjustments in aiming. In particular, probing implicit learning on a small subset of trials tends to overestimate the implicit component (Maresch et al., 2021) as participants may fail to completely dispense with an aiming strategy used in most trials (Avraham et al., 2021).

### Interference as an example of viewing adaptation from the perspective of associative learning

The interference effect we observe in visuomotor adaptation is similar to a well-established phenomenon in associative learning, latent inhibition (Solomon and Moore, 1975). For example, in fear conditioning, pairing a neutral stimulus (conditioned stimulus, CS, e.g., a tone) with an aversive electric shock (unconditioned stimulus, US) leads to a predictive fear response (conditioned response, CR) to the CS. Latent inhibition refers to the finding that conditioning of a CS-US pair is weaker following prolonged exposure to the CS in the absence of the US (CS-alone) (Solomon and Moore, 1975). Thus, if an extended CS-alone block is used to extinguish a conditioned response, reacquisition of this CR will be attenuated when the CS is again paired with the US (Bouton, 1986).

This paradigm is structurally similar to the conditions in which we find attenuation in relearning for sensorimotor adaptation: The pairing of movement with the initial feedback perturbation is broken during the washout block with the introduction of discrepant feedback. Furthermore, both latent inhibition and attenuation upon re-learning in adaptation exhibit context specificity. An attenuated fear response was only evident in the context in which extinction was experienced; when the rats were moved to a different cage between extinction and reacquisition, reacquisition was not attenuated (Bouton, 2002; Bouton and Swartzentruber, 1989). Similarly, adaptation in Experiment 5 was attenuated in a wedge previously associated with a long experience with non-perturbed feedback, but it was spared when tested in a new wedge.

The parallels between these two types of learning may not be incidental (Calame et al., 2023; Raymond et al., 1996; Villalta et al., 2015). Some forms of associative learning, like delay eyeblink conditioning, share important features with implicit sensorimotor adaptation, including strict constraints on the timing of the sensory feedback (Kitazawa et al., 1995; Schneiderman and Gormezano, 1964; Smith et al., 1969; Wang et al., 2024b) and a strong dependency on the integrity of the cerebellum (Donchin et al., 2011; Gao et al., 2016; Izawa et al., 2012; Medina et al., 2000). We have recently considered the utility of a unified framework, viewing implicit sensorimotor adaptation as an associative learning process (Avraham et al., 2022). In a series of experiments, we demonstrated how neutral cues (tone and light) can modulate reach adaptation in a manner that follows the principles of associative learning.

A critical issue in considering a unified framework is to examine the mechanistic overlap between the components of the two processes. We have proposed that the onset of a target, the typical cue for movement initiation in adaptation studies, serves as the CS, one that becomes associated with the perturbed feedback (US). The repeated pairing of the CS and US will increase their associated strength, resulting in an adapted movement (CR) to that target even when the feedback is removed (aftereffect). When viewed in this framework, we see that the target (or the movement plan associated with that target, see Avraham et al., 2022), has CS-like features. For example, adaptation to nearby targets exhibits a generalization pattern similar to that observed in the conditioned eyeblink response to variations in the CS (e.g., tone frequency) (Krakauer et al., 2000; Siegel et al., 1968). Viewing anterograde interference in adaptation as a form of latent inhibition also emphasizes that the target can be viewed as a CS. Repeated movement to a target with veridical feedback will result in attenuated adaptation when the target (CS) is subsequently paired with the perturbed feedback (US); this attenuation is not observed for movements to distant targets.

One problem with the latent inhibition analogy is that we found no attenuation after a long washout block in which feedback was eliminated (Experiment 2) whereas fear conditioning is attenuated following a block of trials in which the CS is presented alone. Eliminating the feedback in sensorimotor adaptation may not be equivalent to eliminating the US in a classical conditioning study, at least when the no-feedback phase is introduced immediately after the learning block. In this condition, the adaptation system may operate as if it is still in the most recent context. As such, when the perturbation is re-introduced, there is no interference. By this hypothesis, we would predict that using an extended block of no-feedback trials at the start of an experiment, where the system is in a non-adapted, baseline state, would result in attenuation in a subsequent learning block.

In summary, a unified framework for associative learning and sensorimotor adaptation helps explain the mechanism underlying memory interference in adaptation. Prolonged experience with non-perturbed feedback (e.g., during baseline or a washout block) produces a strong association, perhaps in a Hebbian-like manner, between the movement plan and that form of feedback (Della-Maggiore et al., 2017). From a Bayesian perspective, the participant forms an inference about the causal structure of the environment, with the non-perturbed feedback being linked to a specific target (the context) (Gershman et al., 2015, 2010). When perturbed feedback is subsequently presented in the same context, the old association interferes with the formation of a new association.

### Relevance of interference to the retention of real-world motor skills

It seems paradoxical that a task as simple as implicit sensorimotor adaptation is so sensitive to interference, even from veridical feedback, whereas complex skills such as bike riding or skiing can be retained over many years, even without practice. However, as shown in the current study, interference requires experience with discrepant feedback within a similar context. The striking retention of complex motor skills may arise because they lack contextual overlap with other behaviors in our motor repertoire. We preserve our ability to ride a bike because we do not perform other skills sufficiently similar to produce interference.

The difference in complexity between simple reaching movements and skilled actions may also limit how well the principles of interference identified in this study extend to more complex movements. We have emphasized that interference is a feature of implicit sensorimotor adaptation, a system designed to ensure that the sensorimotor system remains exquisitely calibrated. While this process is surely operative in the learning and performance of skilled actions, other learning processes will also be relevant to performance; it remains to be seen how they are subject to interference effects. We have noted that explicit strategy use can support savings. However, strategy use could also be prone to interference if a previously learned strategy gets in the way of discovering a new approach (Adamson, 1952). Similarly, other implicit processes essential for highly skilled behavior such as pattern identification and response selection can be subject to pronounced interference effects when the task goal changes (Logan, 1982; Shiffrin and Schneider, 1977).

In the present study, we have used a very restrictive and contrived experimental setup to isolate a particular learning system and characterize one way in which the operation of this system changes as a function of experience. It will be interesting to apply a similar strategy in looking at other learning processes and examining more ecological tasks, with an eye on identifying the conditions that support or interfere with memory consolidation.

## Methods

### Ethics Statement

The study was approved by the Institutional Review Board at the University of California, Berkeley. All participants provided written informed consent to participate in the study and were paid based on a rate of $9.44/hr.

### Participants and experimental setup

236 healthy volunteers (aged 19-38 years; 130 females, 97 males, and 9 identified as ‘other’ or opted not to disclose their sex/gender identity) were tested in six experiments: 44 in each of Experiments 1, 2, 3, and 4, and 60 in Experiment 5. The sample size for each experiment was determined based on the effect size (d=0.6) observed in our previous study showing attenuation of implicit adaptation upon relearning (Avraham et al., 2021), with a significance level of α=0.05 and power of 0.95. The actual number allowed us to counterbalance the direction of the visuomotor perturbation and, where appropriate, the target locations (detailed below).

We used an online platform created within the lab for conducting sensorimotor learning experiments (Tsay et al., 2021a, 2020a). The platform was developed using JavaScript and HTML. Participants were recruited using the crowdsourcing website Prolific (www.prolific.co) and performed the experiment remotely with their personal computers. They were asked to provide information concerning the size of the monitor and the device used (optical mouse or trackpad), as well as their dominant hand (left, right, or ambidextrous). The program was written to adjust the size of the stimuli based on a participant’s monitor size. Due to an error, the data from the questionnaire was not saved. However, our analysis of nearly 2,000 individuals using this platform has shown that implicit visuomotor adaptation does not vary systematically between response devices or handedness (Tsay et al., 2023, 2021a).

### Task

At the beginning of each trial, a white circle (radius: 1% of screen height) appeared at the center of the screen, indicating the start location (Fig. 1A). The participant moved a white cursor (radius: 0.8% of screen height) to the start location. We refer to the body part controlling the cursor — the wrist for mouse users or finger for trackpad users — as the “hand”. After the cursor was maintained in the start location for 500 ms, a blue target (radius: 1% of screen height) appeared at one of four or six locations (see detailed experimental protocols below) positioned along a virtual circle around the start location (radial distance: 25% of screen height).

The participant was instructed to rapidly “reach” towards the target, attempting to slice through the target with their hand. Movement time was calculated as the interval between the time the center of the cursor exceeded the radius of the start location to the time at which the movement amplitude reached the radial distance of the target. To encourage the participants to move fast, the auditory message “too slow” was played if the movement time exceeded 300 ms.

After crossing the target distance, the cursor, when displayed, disappeared and the participant moved back to the start location. The cursor reappeared when the hand was within 25% of the radial distance from the start-to-target distance.

In each experiment, there were three types of feedback conditions (Fig. 1A). On no feedback trials, the cursor was blanked when the hand left the start circle. On veridical feedback trials, the cursor was visible and aligned with the movement direction of the hand; that is, its position corresponded to the standard mapping between the mouse/trackpad and the screen cursor. For the third type of trials, we used task-irrelevant clamped feedback, a visuomotor perturbation that has been shown to drive implicit adaptation with minimal contamination from explicit learning processes (Morehead et al., 2017). Here, the cursor moved along a fixed path, displaced 15° from the target (clockwise or counterclockwise, counterbalanced across participants). We opted to use a 15° clamp as a perturbation of this magnitude is within the range of error sizes (∼6°-60°) shown to induce the maximum rate and extent of adaptation (Kim et al., 2018). The radial position of the cursor corresponded to the participant’s hand position. Thus, the radial position is contingent on the participant’s movement, but the angular position is independent of the participant’s reaching direction. The participant was instructed to ignore the feedback and always reach directly to the target.

To emphasize these instructions, three demonstration trials were performed prior to each clamp block. The target location was fixed for all three trials and chosen to be at least 45° from the two locations on the horizontal meridian (0° and 180°) and 45° from the nearest target used for learning. In all three demonstration trials, 15° clamped feedback was provided. For the first demonstration trial, the participant was instructed to “Reach straight to the left” (180 ), and for the second, “Reach straight to the right” (0 ). Following each of these trials, a message appeared to highlight that the cursor was moving away from their hand and target. For the third demonstration trial, the participant was instructed to “Aim directly for the target”, emphasizing again that they should ignore the cursor.

### Experimental protocol

The main aim of this study was to examine the mechanisms underlying attenuation of implicit sensorimotor adaptation upon relearning. The experiments described below include either one or two blocks with clamped feedback to drive implicit adaptation. The learning blocks were surrounded by blocks with various types of feedback depending on the design of each experiment. Throughout the paper, we use the following convention to describe the experimental protocols. -C refers to a learning block with clamped feedback. The minus (-) sign indicates a perturbation direction that is opposite from the expected direction of adaptation. C (without the minus sign) refers to blocks using a clamp that is rotated in the opposite direction of the clamp used during the learning blocks. V refers to blocks with veridical feedback. N refers to blocks with no feedback. The non-learning blocks, which constitute the main manipulations across our experiments, are highlighted in bold fonts and are separated from the learning blocks using the | sign.

#### Experiment 1: Readaptation following standard washout (-C|CV|-C design)

Experiment 1 was designed to provide a replication of our previous finding that implicit adaptation is attenuated upon relearning (Avraham et al., 2021). The main protocol included two learning blocks separated by a washout block (-C|**CV**|-C). The experimental session consisted of 300 movement cycles, each involving one movement to each of four target locations: 45°, 135°, 225°, 335° (1200 movements in total). The four targets appeared in a pseudorandom and predetermined order within a cycle. The session was composed of the following blocks (Fig. 1B): No Feedback (5 cycles), Veridical Feedback (5 cycles), Learning 1 (80 cycles), Aftereffect 1 (10 cycles), Washout (110 cycles), Learning 2 (80 cycles) and Aftereffect 2 (10 cycles).

The two initial blocks (No Feedback and Veridical Feedback) were included to familiarize the participants with the experimental setup. For these trials, participants were instructed to move their hand directly to the target. By providing veridical feedback in the second block, we expected to minimize idiosyncratic reaching biases. The two learning blocks are the blocks in which we expected to observe implicit adaptation. The aftereffect blocks provided an additional assessment of adaptation. These occurred immediately after the learning blocks, and visual feedback was absent in these trials. Just before the start of the aftereffect block, an instruction screen informed the participant that the cursor would no longer be visible and reminded them that their task was to move directly to the target. The participant pressed a key to start the aftereffect block. While some forgetting might occur during this break, we anticipate that the delay was short. Indeed, the reach angle during the first cycle of the aftereffect block closely matched the reach angle during the last 10 cycles of the learning block ([mean Aftereffect (1st cycle) / Late learning, 95% confidence interval, CI], Learning 1: 0.92, [0.81 1.04], Learning 2: 0.99, [0.85 1.14]).

The washout block was used to bring behavior back to baseline. Typically in studies of relearning in sensorimotor adaptation, washout is administered by removing the perturbation (e.g., the rotation of the cursor) and providing veridical feedback (Albert et al., 2022; Herzfeld et al., 2014; Huang et al., 2011; Krakauer et al., 2005; Morehead et al., 2015; Yin and Wei, 2020; Zarahn et al., 2008). However, assuming adaptation has occurred, the introduction of veridical feedback after adaptation results in a relatively large discrepancy between the expected and observed feedback. We were concerned that this would make participants aware of the change in behavior induced by the clamp, and that this might alter their behavior in the second learning block (e.g., invoke a strategy to offset the anticipated effects of adaptation). To minimize awareness of adaptation, the washout block consisted of two phases (Fig. 1B) (Avraham et al., 2021). In the first phase, we reversed the direction of the clamp. The participant was informed that they would again not have control over the movement direction of the feedback, and they were reminded to ignore the feedback and aim directly for the target. The reversed clamp will reverse adaptation, driving the direction of the hand back towards the target. When the median reach direction was within 5° of the targets for five consecutive cycles, the second phase was implemented with the feedback now veridical.

Demonstration trials were provided at the start of each phase of the washout block, three for the reversed clamp and three for veridical feedback. The demonstration trials were similar to those presented before the learning blocks (same target location and same instructions for where to reach). Note that the demonstration trials for the phase with veridical feedback appeared when the participant’s reaches were relatively close to the target. As such, we expected that participants would be unaware of any residual adaptation at the start of this phase. We did not assess participants’ awareness about their change in behavior during learning. However, when surveyed in a previous study that used the same procedure, none of the participants reported awareness of any learning-related changes in their behavior (Avraham et al., 2021).

We opted to keep the total number of washout cycles fixed. Thus, the number of cycles in each phase of the washout block was determined on an individual basis using the performance criterion described above. All participants experienced at least 70 cycles (mean ± standard deviation, 91.3±13.4) of veridical feedback before the second clamp block, ensuring they had sufficient exposure to unperturbed feedback before the onset of the second learning block.

#### Experiment 2: Readaptation following no feedback washout (-C|N|-C design)

Experiment 2 was designed to test if the attenuation of implicit adaptation upon relearning is due to interference from the feedback experienced during washout. To this end, we tested a condition in which feedback was withheld during the entire washout block (Fig. 2A). The protocol was similar to Experiment 1 with one change: We eliminated the feedback in the 110-cycle washout block (-C|**N**|-C design). As such, the washout block was identical to the preceding aftereffect block.

#### Experiment 3: Readaptation following gradual rotation washout (-C|-RGV|-C design)

Experiment 3 was designed to test if experience with salient, opposite-signed errors at the beginning of the washout block (e.g., the reversed-clamped feedback in Experiment 1) is required to attenuate adaptation upon relearning. The protocol was similar to that used in Experiment 1 with the difference being the procedure used in the washout block. To minimize exposure to large opposite errors, we used rotated feedback that was contingent on the movement of the hand and tailored on an individual basis. For each participant, we calculated the mean reach angle during the first aftereffect block and, for the first cycle of the washout block, rotated the cursor from the actual hand position by that angle multiplied by -1 (Fig. 3A, - C|-**RGV**|-C design). In this way, the rotated visual feedback should counteract the effects of adaptation, with the net effect that the cursor would cross relatively close to the target. On each successive cycle, the size of the rotation was decreased by 1° until it reached zero (veridical feedback). For the remainder of the washout block, the feedback was veridical.

At the beginning of the washout block, participants were told that the cursor would now be aligned with their hand position. We added three demonstration trials to reinforce these instructions. These trials were similar to the demonstration trials used to illustrate the clamped feedback (same target location and instructions of aiming direction). Since the target location used for the demonstration trials (270°) was 45° from the nearest target locations used for adaptation (225° and 315°), we expected a generalization of ∼25% adaptation for reaches to the demonstration locations (Morehead et al., 2017). Thus, the rotation size during these demonstration trials was set to be 25% of the magnitude of the participant’s aftereffect. We assumed this would give the impression that the cursor was aligned with the participant’s movement.

#### Experiment 4: Adaptation following extended veridical feedback (V85|-C)

Experiment 4 was designed to test if extended exposure to veridical feedback was sufficient to produce attenuation of subsequent adaptation. The experimental session consisted of 150 cycles, each consisting of one movement to each of four targets: 30°, 60°, 210°, 240° (Fig. 4A). The four targets appeared in a pseudorandom and predetermined order within a cycle. Adjacent targets were paired [(30°, 60°) and (210°, 240°)] to define a wedge.

For one wedge, we used a -C|**CV**|-C design, consisting of No Feedback (5 cycles), Veridical Feedback (5 cycles), Learning 1 (40 cycles), Aftereffect 1 (10 cycles), Washout (40 cycles), Learning 2 (40 cycles), and Aftereffect 2 (10 cycles). As in Experiment 1, the washout block consisted of a reversed clamp phase and a veridical feedback phase, with the transition between the two phases adjusted on an individual basis (see above). For the second wedge, we used a **V85**|-C design, consisting of No Feedback (5 cycles), Veridical Feedback (85 cycles, with an additional 10 cycle block of No Feedback inserted between cycles 50 and 61), Learning (40 cycles), and Aftereffect (10 cycles). Before each learning block, the participant was provided with a full description of the nature of the feedback for each wedge. In addition, six demonstration trials (three for each wedge) were provided to emphasize the instructions. The association between the wedge location and protocol was counterbalanced across participants.

#### Experiment 5: Adaptation following varying experiences with veridical feedback at distinct contexts (V5|-C / V45|-C / V85|-C design)

Experiment 5 examined if the interference effect from veridical feedback is context specific. The target could appear at one of six locations: 10°, 40°, 130°, 160°, 250°, 280° (Fig. 5A). Pairs of adjacent targets defined three wedges [(10°, 40°), (130°, 160°) and (250°, 280°)], and the number of veridical feedback cycles prior to adaptation varied across the three wedges. The session started with a 5-cycle No Feedback block in which participants moved to each of the six targets, with the locations selected in a random order (within a cycle). This was followed by a Veridical Feedback block (85 cycles). However, the number of wedges included in each cycle changed over the course of the block. In the first phase (40 cycles), the target only appeared in the two locations that defined one wedge; in the second phase (40 cycles), a second wedge was added; and in the third phase (5 cycles), the remaining wedge was included. In this manner, each wedge (counterbalanced across participants) was associated with a different number of veridical feedback trials prior to learning: 5 (**V5**|-C), 45 (**V45**|-C) or 85 (**V85**|-C) cycles. This was followed by a Learning block (40 cycles) and an Aftereffect block (10 cycles); within each of these, all six targets were included in each cycle.

### Data analysis

The kinematic data recorded from the mouse/trackpad were temporarily stored on the Google Firebase database and downloaded for offline analyses using custom-written MATLAB codes. The primary dependent variable was the direction of hand movement (reach angle). Reach angle was defined by two lines, one from the start position to the target and the other from the start position to the hand position at the moment it crossed the radial distance of the target. For participants who experienced a counterclockwise clamp during the learning blocks, the sign of the reach angle was flipped. In this manner, a positive reach angle indicates movement in the opposite direction of the perturbed feedback, the expected change due to adaptation. The reach angle for each movement cycle was calculated by averaging the reach angle of all reaches within a cycle, one to each of the different target locations.

We excluded trials in which the reach angle deviated from the target location by more than 100° and trials in which the absolute trial-to-trial change in reach angle was larger than 20° (Avraham et al., 2021). Based on these criteria, a total of 1.97% of all trials were excluded.

For Experiment 3, we plotted the distribution of errors participants experienced during the non-zero rotation trials of the washout block (Fig. 3B). To evaluate this distribution with respect to an error distribution for veridical feedback (Fig. S2), we used data from another group of participants (N=44). This group experienced an extended, 195-cycle block with veridical feedback. To plot the distribution, we used the data from cycles 100-116, matching the phase of the experiment and the mean number of trials used for calculating the distribution in Experiment 3.

### Statistical analysis

Two statistical approaches were used to analyze the changes in reach angle that occurred in response to the clamped feedback. For the first approach, we used a nonparametric permutation test to identify clusters of cycles in which the reach angle differed between conditions (Avraham et al., 2021; Labruna et al., 2019; Maris and Oostenveld, 2007). For example, to examine within-participant changes in adaptation between the first and the second learning in Experiment 1 (Fig. 1C), a two-tailed paired-sample *t* test was performed for each cycle within the learning and aftereffect blocks.

We defined consecutive cycles in which the difference was significant (p<0.05) as a ‘cluster’ and calculated for each cluster the sum of the t-values that were obtained for the cycles in that cluster (referred to as a t-sum statistic). A null distribution of the t-sum statistic was constructed by performing 10,000 random permutations with the data: For each permutation, the data for a given participant was randomly assigned to “block 1” or “block 2”. For each permuted data set, we performed the same cluster-identification procedure as was done with the actual data and calculated the t-sum statistic for each cluster. In cases where several clusters were found for a given null set permutation, we recorded the t-sum statistic of the cluster with the largest t-sum value. Thus, the generated null distribution is composed of the maximal t-sum values achieved by chance, a method that controls for the multiple comparisons involved in this analysis (Maris and Oostenveld, 2007). Each of the clusters identified in the non-permuted data were considered statistically significant if its t-sum was larger than 95% of the t-sums in the null distribution, corresponding to a *p*-value of 0.05.

The same analysis was used to assess within-participants changes in reach angle between two learning and aftereffect blocks in Experiments 1, 2, 3, and 4. For Experiment 4, we did this procedure three times, for the following comparisons: Learning 2 in the -C|**CV**|-C location vs Learning 2 in the **V85**|-C location, Learning 1 vs Learning 2 in the -C|**CV**|-C location, and Learning 1 in the -C|**CV**|-C location vs Learning 2 in the **V85**|-C location. Therefore, we increased the threshold to identify significant clusters against the null distribution to 98.3%, corresponding to *p*-value of 0.05/3=0.017.

In Experiment 5, we had three conditions that varied in the number of cycles of veridical feedback before learning (85, 45 and 5 cycles). We hypothesized that the extent of adaptation would decrease with the increased experience with veridical feedback. Therefore, we used a linear regression analysis, with the response variable (y) being the mean cycle reach angle and the predictor variable (X) being the number of veridical feedback cycles. We included a constant intercept. We extracted the slope of the regression line for each cycle, and submitted that to the non-parametric, cluster-based permutation analysis. Thus, significant clusters in Figure 5A represent clusters of cycles in which the slope of the regression is significantly different than zero (and negative, representing a decrease in adaptation with increasing experience with veridical feedback) against a null distribution.

The second approach focused on the aftereffect measure of adaptation. This was defined as the mean reach angle over all ten cycles of the aftereffect block. To examine changes in adaptation between the two blocks in Experiments 1, 2, and 3, we used two-tailed paired-sample *t* tests. For the within-participant comparisons of the aftereffect across conditions in Experiments 4, we used a one-way repeated measures ANOVA. The sphericity assumption was violated in Experiment 4, and thus, the *F*-test degrees of freedom was adjusted using the Greenhouse–Geisser correction. For Experiment 5, we used a linear mixed-effect model to fit the aftereffect data, including fixed-effect slope and intercept for the number of veridical cycles before learning (4, 45, and 85) and random-effect slope and intercept for the individual participants. For all of the experiments, the statistical significance threshold was set at the p<0.05. *pB* denotes the Bonferroni corrected p-value. In addition, we report the Bayes factor BF10, the ratio of the likelihood of the alternative hypothesis (H1) over the null hypothesis (H0) (Kass and Raftery, 1995). For *t*-tests, we report the Cohen’s *d* effect size (Cohen, 2013), and for the repeated-measures ANOVAs, we report effect sizes using partial eta-squared (η ^2^). We followed the assumption of normality based on the central limit theorem as the sample size for all experiments was larger than N=30.

### Data and code availability

All raw data files and codes for data analysis are available from the GitHub repository: https://github.com/guyavr/InterferenceRelearningMotorAdaptation.git

## Acknowledgments

We thank Maya Malaviya for her assistance with data collection. This work was supported by grants NS116883, DC077091 from the National Institutes of Health (NIH).

## Competing interests

RBI is a co-founder with equity in Magnetic Tides, Inc.

## Notes

https://github.com/guyavr/InterferenceRelearningMotorAdaptation.git

