## Supplementary Material for "Interference underlies attenuation upon relearning in sensorimotor adaptation"

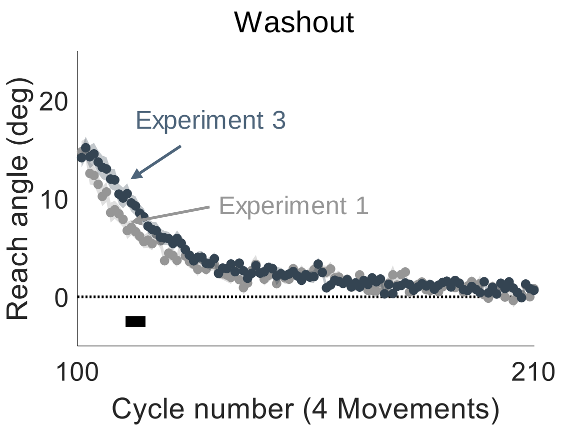


**Fig S1. Early washout in response to a contingent visuomotor rotation that decreases in a gradual manner (Experiment 3) was slower than washout driven by a reversed clamp (Experiment 1).**

Overlaid time courses of mean reach angle during the washout block for Experiment 1 (gray) and Experiment 3 (blue), with the data averaged within each cycle of four movements. The horizontal black bar during the early phase of the washout block denotes a cluster that showed a significant difference between the two experiments, suggesting that the rate of de-adaptation during washout was lower in response to the gradual, contingent rotation schedule that we used compared to a reversed clamp.

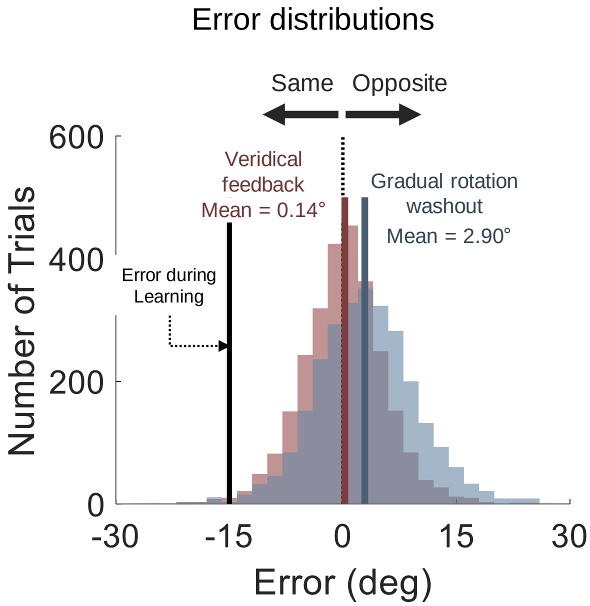


**Fig S2. Comparison of error distributions observed in response to the gradual rotation washout or veridical feedback.**

Overlaid distributions of errors experienced during the non-zero rotation phase of the washout block in Experiment 3 ([mean ± standard deviation], 2.90° ± 10.1°, blue) and with veridical feedback (0.14° ± 10.6°, red). The latter data was obtained from a different group of participants (N=44) which experienced a 195-cycle block with veridical feedback. We used the data from cycles 100-116, matching the phase of the experiment and the mean number of trials used for calculating the distribution in Experiment 3.
